# Integrating three genetic dimensions relating to piglet birth weight: direct and maternal effects on the mean and genetic control of residual variance

**DOI:** 10.1101/2025.06.10.658284

**Authors:** E. Sell-Kubiak, C. Kasper, A. Lepori, J.P. Gutiérrez, N. Formoso-Rafferty, N. Khayatzadeh, I. Cervantes

## Abstract

Uniformity of production traits is desired for different traits in livestock species, including the uniformity of within-litter birth weights in piglets. Birth weight (BW) in pigs is associated with increased vitality and survival until weaning. However, as uniformity of BW increases, the importance of initial weight decreases as competition between piglets decreases. The aim of this study was to estimate the direct and maternal genetic components of BW, jointly with the maternal genetic component of the residual variance for within-litter BW, and their genetic correlations. We used two distinct datasets of Swiss Large White pigs: 1) the experimental farm dataset and 2) the commercial farms dataset, comprising 43,135 and 23,313 records of individual piglet birth weight, respectively. For statistical analysis, the heteroscedastic (or canalising selection) model was used. This model assumes that both the mean BW and the residual variance are affected by systematic and random effects, with the residual variance being heterogeneous and partially under genetic control. Despite the best fitting model was the most complex one including both genetic effects for the mean trait, the results indicated that direct genetic effects, or correlations with such effects, are negligible. The genetic environmental variance for BW ranged between 0.071 and 0.131 for experimental farm and 0.037 to 0.094. The genetic correlation between the mean BW and its variability was always positive and ranged between 0.149 and 0.307 for the experimental farm and between 0.220 and 0.589 for the commercial farms. It is thus sufficient to model BW and its variability by including only the maternal genetic effect for both traits. In addition, even though moderate genetic correlations exist between the mean and the variance of BW, focusing selection on BW uniformity within litters would be preferable to creating a selection index for both traits simultaneously.

**Implications:** Monitoring within-litter birth weight variability is important for efficient piglet production and welfare. Our findings suggest that maternal genetic effects are sufficient to model birth weight and its variability as genetic components of environmental variance. For both traits, direct genetic variance and its correlations with other components are negligible. However, since there are moderate genetic correlations between the mean and variance of birth weight, it is preferable to focus solely on selecting for within-litter birth weight uniformity rather than combining both traits in the selection index. This approach simplifies breeding strategies while maintaining the goal of improving piglets’ welfare.

**Highlights:**

- Data on individual piglet birth weight were collected on Swiss farms over 18 years
- Heteroscedastic animal models integrated three dimensions of genetic components
- The direct genetic effect and its correlations are minimal and irrelevant for selection
- Maternal genetic effects are crucial for birth weight and its variability
- Selection for uniformity is preferable to selection for both traits simultaneously

## Introduction

Uniformity is an important quality of various economically significant traits in livestock production, one of which is piglet birth weight. Birth weight (BW) in pigs is associated with increased vitality (Rutherford et al., 2013) and survival until weaning (Kapell et al., 2011; Knol et al., 2010). However, as the within-litter BW becomes more uniform, the importance of initial weight diminishes as competition between piglets decreases (Andersen et al., 2011; Canario et al., 2010). To avoid issues related to a high number of piglets nursed by one sow and to obtain a homogenized BW across piglets, cross-fostering is used (Baxter et al., 2013). This practice increases preweaning survival and, therefore, farm productivity (Putz et al., 2015); however, at the same time, it increases labour costs and poses immunity concerns (Baxter et al., 2013).

Selection for increased uniformity of within-litter BW could be implemented to avoid intervention practices such as cross-fostering. For this purpose, estimating variance components of BW variability is a priority. BW variability can be studied by the heterogeneity of residual variance of the trait of interest, which was previously presented in pigs by Sell-Kubiak et al. (Sell-Kubiak et al., 2015; Sell-Kubiak et al., 2015) and in mice by Formoso-Rafferty et al. (Formoso-Rafferty et al., 2016). In addition, in their experiment on mice, Formoso-Rafferty et al. (Formoso-Rafferty et al., 2019, 2022) reported that selecting animals with reduced environmental BW variability is possible, which also led to more robust and feed-efficient animals. Thus, selection for uniformity in within-litter BW could lead to more ethical and efficient livestock production because it results in litters that are easier to manage (Rutherford et al., 2013) and piglets that are more likely to survive until weaning (Damgaard et al., 2013). Furthermore, the application of a canalising (SanCristobal-Gaudy et al., 1998) selection model provides a framework to better understand and model the genetic and environmental factors underlying BW variability. This model allows the residual variance to vary across individuals depending on genetic and environmental influences and includes a specific genetic effect on variability, thereby accounting for the model’s heterogeneous residual variance. Notably, BW and its variability are considered maternal traits (Sell-Kubiak et al., 2015; Damgaard et al., 2013; Nielsen et al., 2013; Canario et al., 2010), and sows differ from each other in their sensitivity to environmental disturbances (Rönnegård et al., 2010). Although successful selection for environmental BW variability can be performed by assuming that these traits are maternal (Formoso-Rafferty et al., 2016), there is evidence for a direct genetic component in the mean BW (Alves et al., 2018; Kaufmann et al., 2000).

Thus, this study aimed to estimate the direct genetic and maternal genetic components of BW jointly with the maternal genetic component of the residual variance for within-litter BW and their genetic correlations in two datasets collected from two types of farms in Switzerland.

## Methods

### Animals

The study was performed in two different locations, yielding two distinct datasets: 1) the Swiss experimental farm dataset comprising 43,135 records of BW from 3,163 litters of 986 sows collected between 2004 and 2022 and pedigree data for 45,737 individuals; and 2) the Swiss commercial farms dataset comprising 23,313 BW records from 1,748 litters of 813 sows from two farms collected between 2004 and 2022 at two different farms and 27,285 individuals in the pedigree. The descriptive statistics of both datasets are presented in Tables 1 and 2. In Switzerland, free farrowing has been required by law since 2007 (Swiss Animal Protection Ordinance). This means that farrowing pens must be designed such that the sow can turn freely, and thus, farrowing crates are not allowed.

**Table 1.** Number of records, mean and standard deviation, minimum and maximum for the analysed traits in the experimental farm dataset.

|  | **BW mean**<br>**(kg)** | **Standard**<br>**Deviation** | **Minimum** | **Maximum** | **Number of**<br>**records** |
| --- | --- | --- | --- | --- | --- |
| **All** | 1.42 | 0.38 | 0.24 | 3.00 | 43135 |
| **Sex** |  |  |  |  |  |
| Male | 1.45 | 0.39 | 0.30 | 3.00 | 22522 |
| Female | 1.40 | 0.37 | 0.24 | 2.93 | 20571 |
| Hermaphrodites | 1.28 | 0.28 | 0.79 | 1.66 | 11 |
| Unknown | 1.26 | 0.35 | 0.45 | 2.20 | 31 |
| **Litter size** |  |  |  |  |  |
| 1 (<7) | 1.82 | 0.40 | 0.30 | 2.80 | 762 |
| 2 (7) | 1.75 | 0.37 | 0.54 | 2.56 | 511 |
| 3 (8) | 1.61 | 0.44 | 0.43 | 2.80 | 616 |
| 4 (9) | 1.67 | 0.39 | 0.39 | 2.78 | 1008 |
| 5 (10) | 1.57 | 0.37 | 0.26 | 2.65 | 1613 |
| 6 (11) | 1.60 | 0.38 | 0.41 | 2.79 | 2332 |
| 7 (12) | 1.50 | 0.36 | 0.39 | 2.73 | 3289 |
| 8 (13) | 1.49 | 0.37 | 0.40 | 2.93 | 4205 |
| 9 (14) | 1.45 | 0.35 | 0.32 | 2.62 | 5006 |
| 10 (15) | 1.39 | 0.36 | 0.32 | 2.76 | 5188 |
| 11 (16) | 1.38 | 0.36 | 0.32 | 3.00 | 5563 |
| 12 (17) | 1.32 | 0.34 | 0.31 | 2.69 | 4573 |
| 13 (18) | 1.32 | 0.35 | 0.28 | 2.39 | 3222 |
| 14 (19) | 1.27 | 0.35 | 0.37 | 2.38 | 2488 |
| 15 (20) | 1.25 | 0.35 | 0.24 | 2.50 | 1344 |
| 16 (21) | 1.24 | 0.34 | 0.31 | 2.26 | 756 |
| 17 (22-27) | 1.17 | 0.34 | 0.35 | 2.20 | 659 |
| **Number of**<br>**parturitions** |  |  |  |  |  |
| 1 | 1.34 | 0.34 | 0.26 | 2.73 | 12624 |
| 2 | 1.49 | 0.38 | 0.31 | 3.00 | 9271 |
| 3 | 1.47 | 0.40 | 0.32 | 2.76 | 7065 |
| 4 | 1.45 | 0.39 | 0.24 | 2.73 | 5376 |
| 5 | 1.44 | 0.38 | 0.40 | 2.79 | 3885 |
| 6 | 1.41 | 0.39 | 0.30 | 2.44 | 2504 |
| 7 | 1.40 | 0.38 | 0.43 | 2.41 | 1334 |
| 8 | 1.38 | 0.41 | 0.35 | 2.51 | 631 |
| 9 | 1.41 | 0.43 | 0.42 | 2.63 | 299 |
| 10 (>9) | 1.46 | 0.42 | 0.59 | 2.49 | 146 |

**Table 2.** Number of records, mean and standard deviation, minimum and maximum for the analysed traits in the commercial farms dataset.

|  | **BW mean**<br>**(kg)** | **Standard**<br>**Deviation** | **Minimum** | **Maximum** | **Number of**<br>**records** |
| --- | --- | --- | --- | --- | --- |
| **All** | 1.55 | 0.36 | 0.40 | 2.70 | 23313 |
| **Breed** |  |  |  |  |  |
| Breed 1 | 1.52 | 0.35 | 0.40 | 2.66 | 12530 |
| Breed 51 | 1.50 | 0.31 | 0.61 | 2.65 | 755 |
| Breed 99 | 1.48 | 0.31 | 0.43 | 2.28 | 1014 |
| Breed 104 | 1.61 | 0.37 | 0.41 | 2.70 | 9014 |
| **Sex** |  |  |  |  |  |
| Male | 1.57 | 0.36 | 0.43 | 2.70 | 12016 |
| Female | 1.53 | 0.35 | 0.40 | 2.67 | 11297 |
| **Litter Size** |  |  |  |  |  |
| 1 (<6) | 1.79 | 0.33 | 0.83 | 2.55 | 172 |
| 2 (6) | 1.85 | 0.35 | 0.94 | 2.65 | 168 |
| 3 (7) | 1.77 | 0.37 | 0.60 | 2.58 | 239 |
| 4 (8) | 1.75 | 0.32 | 0.80 | 2.70 | 419 |
| 5 (9) | 1.72 | 0.35 | 0.68 | 2.65 | 669 |
| 6 (10) | 1.70 | 0.35 | 0.43 | 2.67 | 1229 |
| 7 (11) | 1.63 | 0.35 | 0.47 | 2.66 | 1874 |
| 8 (12) | 1.63 | 0.33 | 0.48 | 2.55 | 2260 |
| 9 (13) | 1.57 | 0.34 | 0.45 | 2.65 | 2928 |
| 10 (14) | 1.52 | 0.34 | 0.44 | 2.57 | 3216 |
| 11 (15) | 1.52 | 0.35 | 0.43 | 2.61 | 2734 |
| 12 (16) | 1.48 | 0.35 | 0.40 | 2.52 | 2775 |
| 13 (17) | 1.48 | 0.35 | 0.50 | 2.55 | 1935 |
| 14 (18) | 1.43 | 0.33 | 0.42 | 2.47 | 1145 |
| 15 (19) | 1.45 | 0.34 | 0.53 | 2.63 | 746 |
| 16 (20-25) | 1.34 | 0.34 | 0.46 | 2.50 | 804 |

The experimental farm at Agroscope Posieux maintains the dam line of the Swiss Large White breed and has a self-replacement system, recruiting replacement gilts exclusively from its own breeding herd. Thus, all sows in this dataset were from the Large White dam line. Sows were kept in groups of 8 to 14 animals in pens of approximately 43 m^2^. The sows in the same group were inseminated and gave birth at the same time (farrowing batches). Between the 110th day of gestation and the 26th day of lactation, the sows were housed in individual 7.1 m^2^ pens (5.9 m^2^ solid concrete floor and 1.2 m^2^ slatted concrete floor). The piglets had exclusive access to a separate, covered area (0.8 m^2^) heated with infrared lamps, which was not accessible to the sow. The pen was equipped with an automatic feeder (Schauer Spotmix, Agrotonic GmbH, Prambachkirchen, Austria) allowing the provision of predefined amounts of feed. Piglets were individually weighed within 24 hours of farrowing by trained piggery staff via scales with 5 g precision. Before weighing, all piglets in one litter were placed in a box warmed by an infrared lamp, and each piglet was then removed individually for measurement. The scales automatically accounted for movement by recording multiple weight measurements over a set period and averaging them.

Two nucleus farms from SUISAG were also included in this study. Both farms inseminated Large White sows with Large White (sire and dam lines) and Landrace boars. The litters were primarily purebred Large White, F1 crosses, a small amount of crossbred Large White, and the other unknown crosses. Sows were kept in groups of 7 to 14 animals in waiting pens before being transferred to farrowing pens on day 110 of gestation. Sows were kept in the farrowing pens for 26-30 days of lactation. Each farrowing pen provided 0.8 m^2^ of space for the born piglets and was equipped with infrared heating lamps. After 24h from birth, the piglets born in a litter were placed in the scale box together (Mettler Toldeo with a precision of 5g). Each piglet was then removed one by one, and the decrease in weight was recorded as the individual birth weight. Stillborn piglets were not included in the measurement.

### Statistical analysis

The systematic effects used in the models and curated in the datasets were the mother’s age: from 300 to 1956 days (commercial farm) or the number of parturitions: 10 levels (experimental farm), sex, two levels for commercial farm and 4 for the experimental farm (male, female, hermaphrodites and unknown), litter size including 16 levels (1-5, 6, 7… 20-25) for commercial farm and 17 levels (1-6, 7… 20, 21, 22-27) for the experimental farm, a comparison group that included combinations of farm-month years for commercial farm (75 levels) and 357 batches for the experimental farm and the male breed, with four levels (commercial farm). The litter effect was included as a random effect and included 1755 and 3165 levels for commercial and experimental farms, respectively.

### Homoscedastic Model

In the first step, a classical animal model assuming homoscedastic residual variance was implemented. This model is referred to as HO:

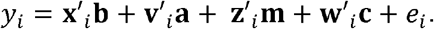

In this model, *y*_*i*_ is the individual birth weight of animal *i*; **b** are the vectors of the systematic effects; **a** is the vector of the direct genetic effect (individual); **m** is the vector of the genetic effect of the mother of individual i (maternal genetic effect); **c** is a vector of the litter effect; *e*_*i*_ is the residual; and *x*′_*i*_, *z*′_*i*_, and *w*′_*i*_ are the incidence matrices for systematic, litter and animal effects, respectively.

### Heteroscedastic Model

The heteroscedastic (or canalising selection) model assumes that both the mean birth weight level and the residual variance are affected by systematic and random effects, with the residual variance being heterogeneous and partially under genetic control.

The more complex heteroscedastic model included the direct genetic effect (individual) and the maternal genetic effect for BW and the maternal genetic effect for its variability:

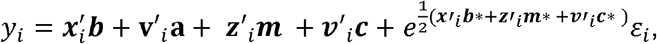

where *y*_*i*_ is the individual birth weight of animal i; * indicates the parameters associated with the environmental variance, i.e., the variability of BW; **b** and **b\*** are the vectors of the systematic effects; **a** is the vector of the additive genetic effect (individual); **m** and **m\*** are the vectors of the genetic effect of the mother of individual i; **c** and **c\*** are the vectors of the litter effect; ***x***′_*i*_, ***z***′_*i*_, and ***w***′*i*are the incidence matrices for systematic, litter and animal effects, respectively; and ε_*i*_ is the standardized error. The genetic effects **a, m** and **m\*** are jointly distributed and are assumed to be Gaussian, as follows in the maternal model:

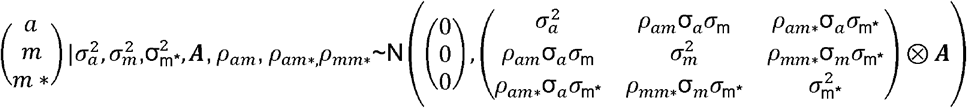

where **A** is the additive genetic relationship matrix; 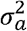 is the direct additive genetic variance of the trait; 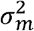 is the maternal additive genetic variance of the trait; 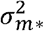 is the maternal additive genetic variance affecting the environmental variance of the trait; ρ_am_, ρ_am_^*^ and ρmm^*^ are the coefficients of genetic correlations between them; and ⊗ denotes the Kronecker product.

The litter effects c and c* are jointly distributed and are assumed to be Gaussian, as follows:

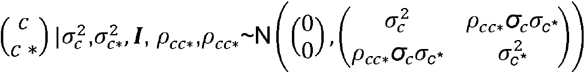

The heteroscedastic models (Table 3), referred to as HE, included different combinations of random effects and correlations between them:

**Table 3.** Different variance components included in the tested models (HO, homoscedastic model; HE, heteroscedastic model).

| Model | Var(a) | Var(m) | Var(m*) | Var(c) | Var(c*) | Var(e) | Cov(am) | Cov(am*) | pmm* | pam | pam* | pcc* |
| --- | --- | --- | --- | --- | --- | --- | --- | --- | --- | --- | --- | --- |
| HO | X | X |  | X |  | X | X |  |  | X |  |  |
| HE_1 |  | X | X | X | X |  |  |  | X |  |  |  |
| HE_2 |  | X | X | X | X |  |  |  |  |  |  | X |
| HE_3 | X | X | X |  | X |  | X | X | X | X | X |  |
| HE_4 | X | X | X | X |  |  | X | X | X | X | X |  |
| HE_5 | X | X | X | X | X |  | X | X | X | X | X | X |
| HE_6 | X |  | X | X | X |  |  | X |  | X |  | X |
| HE_7 | X | X | X | X | X |  | X | X | X | X | X |  |
| HE_8 | X | X | X |  |  |  | X | X | X | X | X |  |

- Model HE_1 included the maternal genetic effect for the mean trait (m) and its variability (m*), their genetic correlation, ρmm^*^, and the litter effect for the mean trait and its variability (c, c*) without correlation between them.
- Model HE_2 included the same effects as HE_1 and the correlation between both litter effects (ρcc^*^).
- Model HE_3 included the direct genetic effect for the mean trait (a), the maternal genetic effect for the mean trait (m) and its variability (m*), and the genetic correlation between them, ρ_am_, ρ_am_^*^ and ρmm^*^. The litter effect was only fitted for the mean trait (c).
- Model HE_4 included the direct genetic effect for the mean trait (a), the maternal genetic effect for the mean trait (m) and its variability (m*), and the genetic correlation between them, ρ_am_, ρ_am_^*^ and ρmm^*^. The litter effect was only fitted for BW variability (c*).
- Model HE_5 was the most complex model, including the direct genetic effect for the mean trait (a), the maternal genetic effect for the mean trait (m) and its variability (m*), and the genetic correlation between them, ρ_am_, ρ_am_^*^ and ρmm^*^. The litter effect was fitted for the mean trait (c) and its variability (c*). Additionally, the correlation between litter effects was included (ρcc^*^).
- Model HE_6 included the same effects as HE_5, except for the maternal genetic effect for the mean trait (m*) and the associated genetic correlations.
- Model HE_7 was the same as HE_5 but without a correlation between the two litter effects.
- Model HE_8 was the same as HE_5 but had no effect on the mean trait or its variability.

The Akaike information criterion (AIC) and Bayesian information criterion (BIC) were used to compare the models and select those that best fit the data.

Since the HE models’ residual variance is heterogeneous by default, a residual variance can also be estimated for a particular level of systematic effects. A global residual variance was estimated in the HE models by adding the averages of the estimates of all the levels within systematic effects. To maintain the estimability of the corresponding linear combination, the solutions for all levels of each of the other systematic effects were averaged and added to the solution for a particular desired level of the systematic effect. The global heritability (h^2^) for the mean trait and each level of systematic effect was subsequently computed (Formoso-Rafferty et al., 2017). ASReml 4.2 (Gilmour et al., 2015) was used to estimate the variance components for the heteroscedastic model and homoscedastic model. GSEVM (Ibáñez-Escriche et al., 2010) software was used to obtain solutions for systematic effects and BW heritabilities via the HE_1 model. Note: The HE_1 model was selected since the GSEVM does not allow for the inclusion of more than one additive genetic effect.

## Results

### Systematic effects for BW in the models

Figure 1 shows the magnitude of the systematic effects on BW via the HE_1 model in the experimental farm dataset. Males were heavier than females were; hermaphrodites were lighter than females were. The BW decreased with litter size. It increased considerably from the first to the second parturition, remained similar to the second at the third parturition, and then gradually decreased in subsequent parturitions. Similar results were found on the commercial farms (Figure 2). Here, the sow’s age was positively related to BW, i.e., older sows produced piglets with higher BW. We also found differences in BW depending on the breed on the commercial farm, with one paternal breed resulting in slightly heavier piglets than the other three breeds.

**Figure 1.**
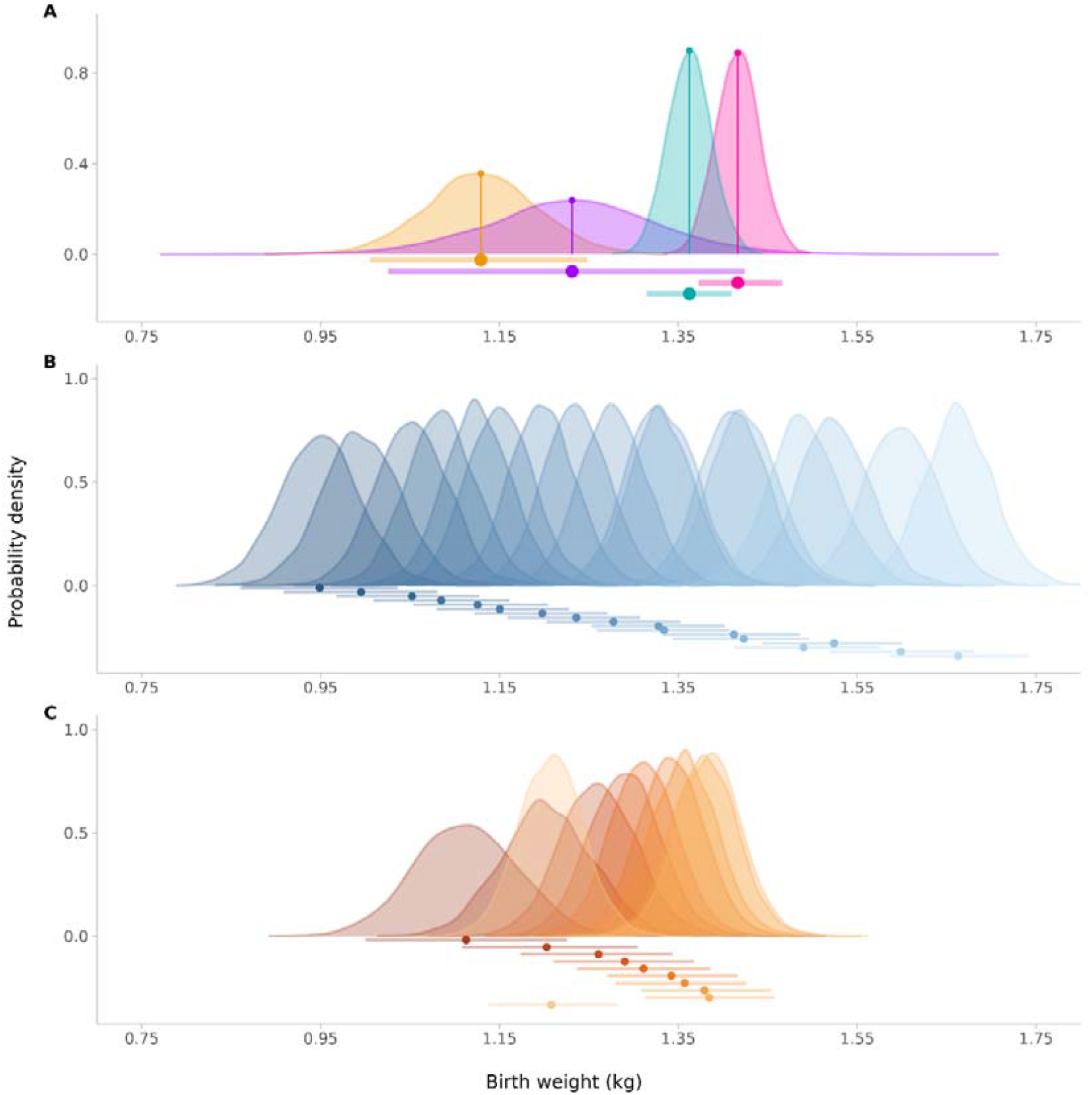
Probability density of the posterior distributions of systematic effects solutions of sex (A), litter size (B), and number of parturitions (C) on birth weight in the experimental farm dataset. A: turquoise - female, pink - male, purple - hermaphrodite, orange - sex unknown. B and C: darker colours indicate larger litter sizes and greater numbers of parturitions, respectively.

**Figure 2.**
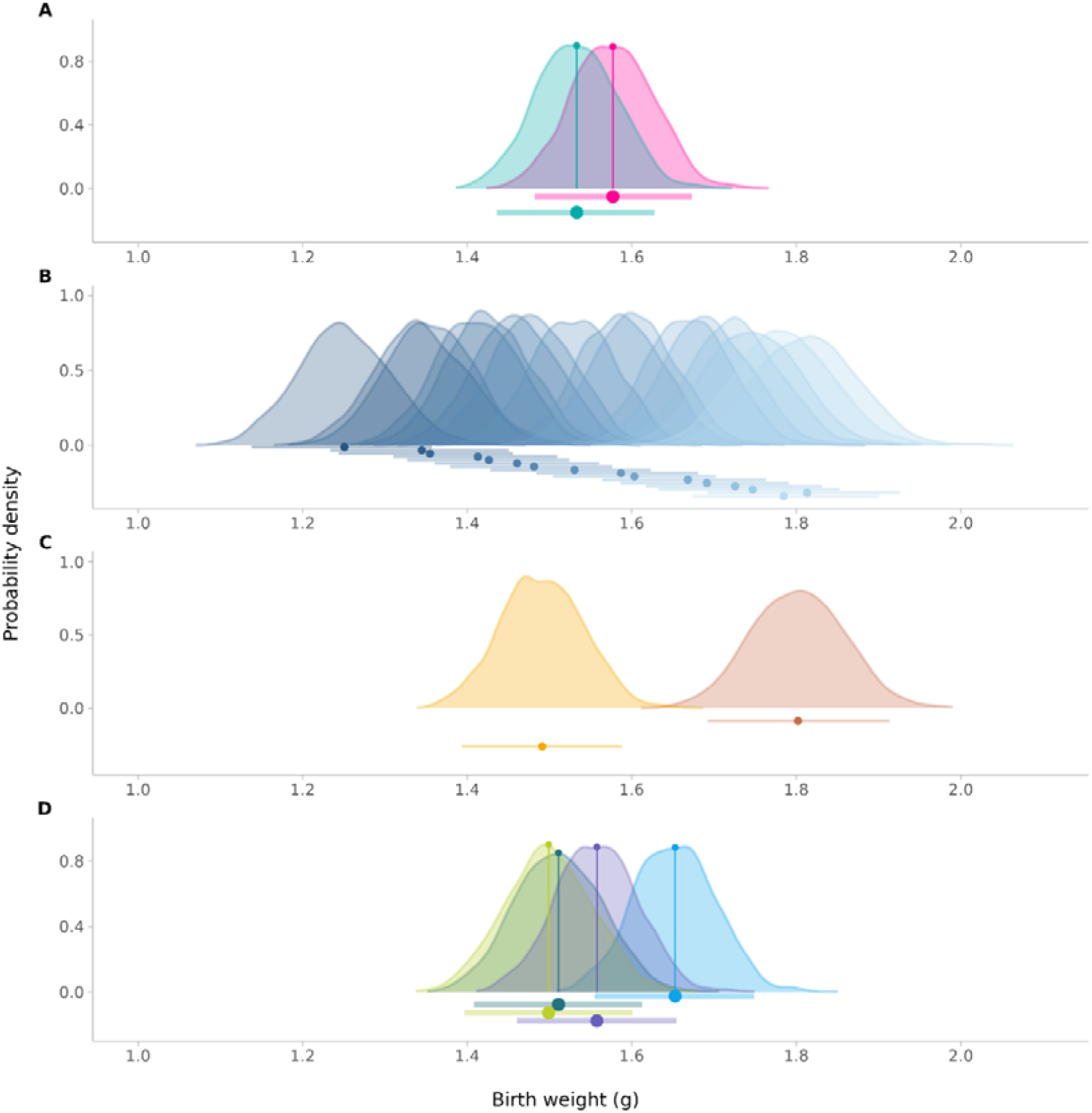
Probability density of the posterior distributions of systematic effects solutions of sex, litter size, breed, and age on birth weight in the commercial farm dataset. A: sex; turquoise - female, pink - male. B: litter size; darker colours indicate larger litter sizes. C: sow’s age; darker colours indicate greater age. D: sires’ breed.

### Estimates of variance components and heritabilities

Tables 4 and 5 show the estimates of variance components and correlations between them in the different models applied to both datasets. The results of the HO model indicated that the maternal genetic variance was greater (0.029 and 0.023 in the experimental and commercial farms, respectively) than the individual genetic variance was (0.013 and 0.005 in the experimental and commercial farms, respectively). Heritabilities for BW were 0.095 (±0.0194) for the direct genetic component and 0.217 (±0.0221) for the maternal effect at the experimental farm. In contrast, lower values were found at the commercial farm, with values of 0.038 (±0.0137) and 0.195 (±0.0261) for direct and maternal effects, respectively. The HE_1 model (including maternal effects only) yielded a genetic environmental variance for BW of 0.075 in the experimental farm data compared with 0.037 in the commercial farm data. The individual genetic variance for BW was close to 0 when the maternal genetic effect was added to the HE model. When the maternal effect was excluded, the variance explained was not absorbed by the direct genetic effect (Model HE_6), even though the variance of this effect slightly increased. The magnitude of the litter effect was greater for BW variation than for individual BW.

**Table 4.**
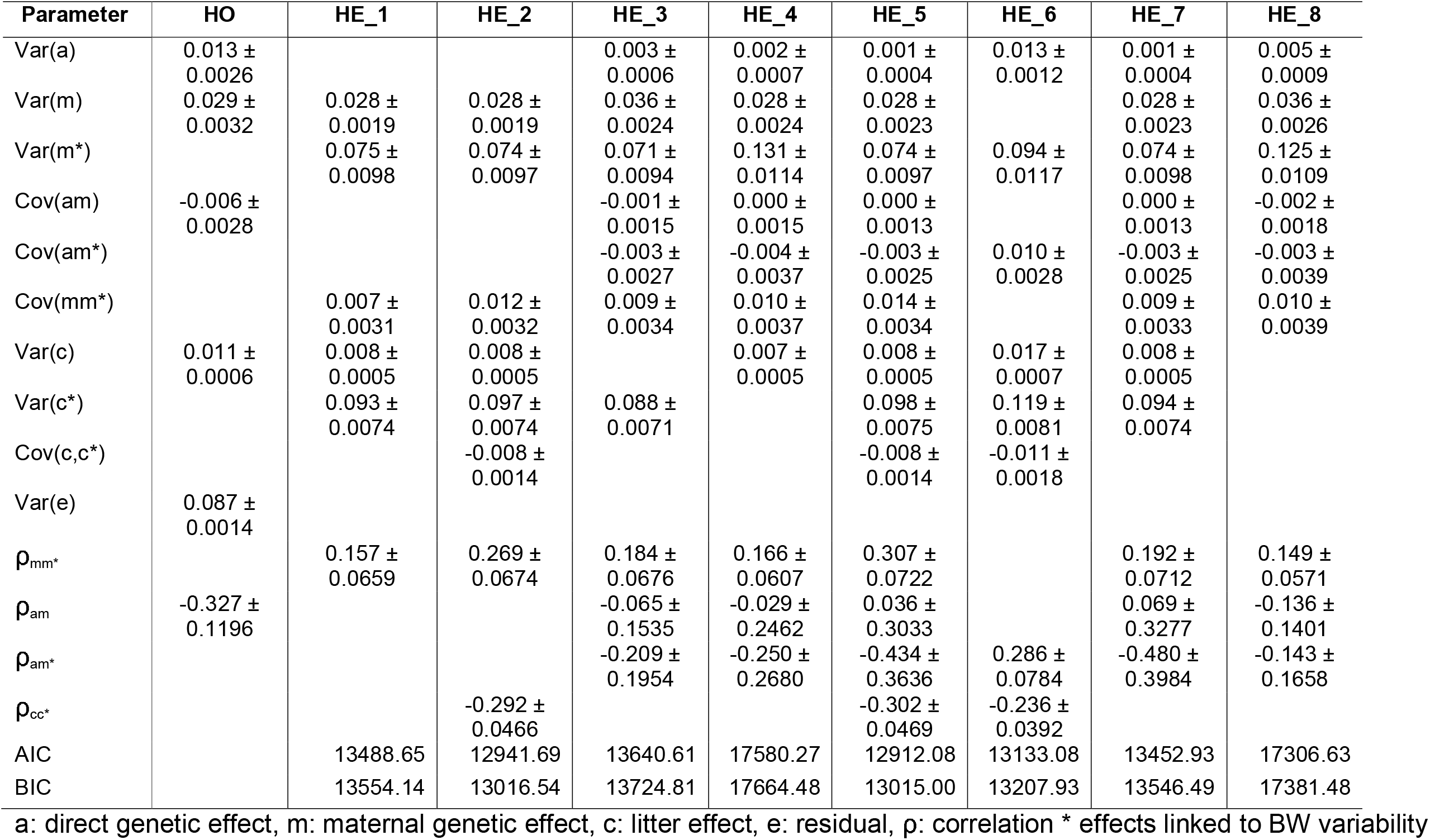
Estimates (mean ± SE) of variance components and correlations in the different models for piglet birth weight applied to the experimental farm dataset (HO, homoscedastic model; HE, heteroscedastic model).

| Parameter | HO | HE_1 | HE_2 | HE_3 | HE_4 | HE_5 | HE_6 | HE_7 | HE_8 |
| --- | --- | --- | --- | --- | --- | --- | --- | --- | --- |
| Var(a) | 0.013 ±<br>0.0026 | | | 0.003 ±<br>0.0006 | 0.002 ±<br>0.0007 | 0.001 ±<br>0.0004 | 0.013 ±<br>0.0012 | 0.001 ±<br>0.0004 | 0.005 ±<br>0.0009 |
| Var(m) | 0.029 ±<br>0.0032 | 0.028 ±<br>0.0019 | 0.028 ±<br>0.0019 | 0.036 ±<br>0.0024 | 0.028 ±<br>0.0024 | 0.028 ±<br>0.0023 | | 0.028 ±<br>0.0023 | 0.036 ±<br>0.0026 |
| Var(m*) | | 0.075 ±<br>0.0098 | 0.074 ±<br>0.0097 | 0.071 ±<br>0.0094 | 0.131 ±<br>0.0114 | 0.074 ±<br>0.0097 | 0.094 ±<br>0.0117 | 0.074 ±<br>0.0098 | 0.125 ±<br>0.0109 |
| Cov(am) | -0.006 ±<br>0.0028 | | | -0.001 ±<br>0.0015 | 0.000 ±<br>0.0015 | 0.000 ±<br>0.0013 | | 0.000 ±<br>0.0013 | -0.002 ±<br>0.0018 |
| Cov(am*) | | | | -0.003 ±<br>0.0027 | -0.004 ±<br>0.0037 | -0.003 ±<br>0.0025 | 0.010 ±<br>0.0028 | -0.003 ±<br>0.0025 | -0.003 ±<br>0.0039 |
| Cov(mm*) | | 0.007 ±<br>0.0031 | 0.012 ±<br>0.0032 | 0.009 ±<br>0.0034 | 0.010 ±<br>0.0037 | 0.014 ±<br>0.0034 | | 0.009 ±<br>0.0033 | 0.010 ±<br>0.0039 |
| Var(c) | 0.011 ±<br>0.0006 | 0.008 ±<br>0.0005 | 0.008 ±<br>0.0005 | | 0.007 ±<br>0.0005 | 0.008 ±<br>0.0005 | 0.017 ±<br>0.0007 | 0.008 ±<br>0.0005 | |
| Var(c*) | | 0.093 ±<br>0.0074 | 0.097 ±<br>0.0074 | 0.088 ±<br>0.0071 | | 0.098 ±<br>0.0075 | 0.119 ±<br>0.0081 | 0.094 ±<br>0.0074 | |
| Cov(c,c*) | | | -0.008 ±<br>0.0014 | | | -0.008 ±<br>0.0014 | -0.011 ±<br>0.0018 | | |
| Var(e) | 0.087 ±<br>0.0014 | | | | | | | | |
| $\rho_{mm^*}$ | | 0.157 ±<br>0.0659 | 0.269 ±<br>0.0674 | 0.184 ±<br>0.0676 | 0.166 ±<br>0.0607 | 0.307 ±<br>0.0722 | | 0.192 ±<br>0.0712 | 0.149 ±<br>0.0571 |
| $\rho_{am}$ | -0.327 ±<br>0.1196 | | | -0.065 ±<br>0.1535 | -0.029 ±<br>0.2462 | 0.036 ±<br>0.3033 | | 0.069 ±<br>0.3277 | -0.136 ±<br>0.1401 |
| $\rho_{am^*}$ | | | | -0.209 ±<br>0.1954 | -0.250 ±<br>0.2680 | -0.434 ±<br>0.3636 | 0.286 ±<br>0.0784 | -0.480 ±<br>0.3984 | -0.143 ±<br>0.1658 |
| $\rho_{cc^*}$ | | | -0.292 ±<br>0.0466 | | | -0.302 ±<br>0.0469 | -0.236 ±<br>0.0392 | | |
| AIC |  | 13488.65 | 12941.69 | 13640.61 | 17580.27 | 12912.08 | 13133.08 | 13452.93 | 17306.63 |
| BIC |  | 13554.14 | 13016.54 | 13724.81 | 17664.48 | 13015.00 | 13207.93 | 13546.49 | 17381.48 |

a: direct genetic effect, m: maternal genetic effect, c: litter effect, e: residual, $\rho$ : correlation \* effects linked to BW variability

**Table 5.** Estimates (mean ± SE) of variance components and correlations in the different models for piglet birth weight applied to the commercial farm dataset (HO, homoscedastic model; HE, heteroscedastic model).

|  | HO | HE_1 | HE_2 | HE_3 | HE_4 | HE_5 | HE_6 | HE_7 | HE_8 |
| --- | --- | --- | --- | --- | --- | --- | --- | --- | --- |
| Var(a) | 0.005 ±<br>0.0017 | | | 0.005 ±<br>0.0009 | 0.002 ±<br>0.0010 | 0.000 ±<br>0.0007 | 0.005 ±<br>0.0012 | 0.000 ±<br>0.0007 | 0.008 ±<br>0.0014 |
| Var ± m) | 0.023 ±<br>0.0033 | 0.023 ±<br>0.0022 | 0.023 ±<br>0.0022 | 0.040 ±<br>0.0033 | 0.022 ±<br>0.0030 | 0.023 ±<br>0.0028 | | 0.022 ±<br>0.0028 | 0.040 ±<br>0.0036 |
| Var ± m*) | | 0.037 ±<br>0.0094 | 0.040 ±<br>0.0095 | 0.044 ±<br>0.0099 | 0.094 ±<br>0.0115 | 0.067 ±<br>0.0123 | 0.042 ±<br>0.0101 | 0.037 ±<br>0.0094 | 0.099 ±<br>0.0117 |
| Cov ± am) | -0.001 ±<br>0.0033 | | | -0.001 ±<br>0.0023 | 0.000 ±<br>0.0023 | 0.001 ±<br>0.0019 | | 0.001 ±<br>0.0020 | -0.002 ±<br>0.0028 |
| Cov ±<br>am*) | | | | -0.002 ±<br>0.0035 | -0.004 ±<br>0.0049 | -0.002 ±<br>0.0032 | 0.008 ±<br>0.0026 | -0.001 ±<br>0.0032 | -0.004 ±<br>0.0053 |
| Cov ±<br>mm*) | | 0.012 ±<br>0.0032 | 0.017 ±<br>0.0034 | 0.015 ±<br>0.0042 | | 0.018 ±<br>0.0037 | | 0.012 ±<br>0.0035 | 0.014 ±<br>0.0047 |
| Var ± c) | 0.010 ±<br>0.0008 | 0.013 ±<br>0.0009 | 0.013 ±<br>0.0009 | | 0.012 ±<br>0.0089 | 0.017 ±<br>0.0009 | 0.025 ±<br>0.0012 | 0.013 ±<br>0.0009 | |
| Var ± c*) | | 0.119 ±<br>0.0112 | 0.122 ±<br>0.0112 | 0.105 ±<br>0.0108 | | 0.123 ±<br>0.0112 | 0.131 ±<br>0.0117 | 0.120 ±<br>0.0113 | |
| Cov ±<br>c,c*) | | | -0.009 ±<br>0.0023 | | | -0.009 ±<br>0.0023 | -0.006 ±<br>0.0027 | | |
| Var ± e) | 0.083 ±<br>0.0011 | | | | | | | | |
| $\rho_{mm^*}$ | | 0.403 ±<br>0.1059 | 0.556 ±<br>0.1054 | 0.366 ±<br>0.0979 | 0.295 ±<br>0.0869 | 0.589 ±<br>0.1159 | | 0.429 ±<br>0.1181 | 0.220 ±<br>0.0731 |
| $\rho_{am}$ | -0.117 ±<br>0.2561 | | | -0.073 ±<br>0.1593 | 0.042 ±<br>0.3406 | 0.209 ±<br>0.7703 | | 0.325 ±<br>1.0189 | -0.114 ±<br>0.1465 |
| $\rho_{am^*}$ | | | | -0.107 ±<br>0.2337 | -0.263 ±<br>0.3568 | -0.406 ±<br>0.9604 | 0.536 ±<br>0.1677 | -0.435 ±<br>1.2269 | -0.153 ±<br>0.1813 |
| $\rho_{cc^*}$ | | | -0.228 ±<br>0.0562 | | | -0.232 ±<br>0.0565 | -0.096<br>± 0.0479 | | |
| AIC |  | 6648.11 | 6444.69 | 7084.36 | 9852.27 | 6441.98 | 6706.46 | 6716.41 | 9605.62 |
| BIC |  | 6709.33 | 6514.66 | 7163.07 | 9930.99 | 6538.19 | 6776.43 | 6803.86 | 9675.58 |

a: direct genetic effect, m: maternal genetic effect, c: litter effect, e: residual, p: correlation \* effects linked to BW variability

Figures 3 and 4 show the heritabilities for BW depending on the level of systematic effect using the HE_1 model in the experimental farm and commercial farm datasets. At the experimental farm, females presented greater heritability than males did; for litter size, intermediate values were less consistent, and the BW variation increased with parity number. With respect to the commercial farm, female piglets were more consistent in BW despite having lower weights, and litter sizes greater than nine piglets had greater variability than smaller litters did. The age of the sow positively affected the mean BW but increased the degree of BW variation. The expected global maternal heritability for BW was 0.208 (0.0205) for the commercial farm and 0.214 (0.0289) for the experimental farm.

**Figure 3.**
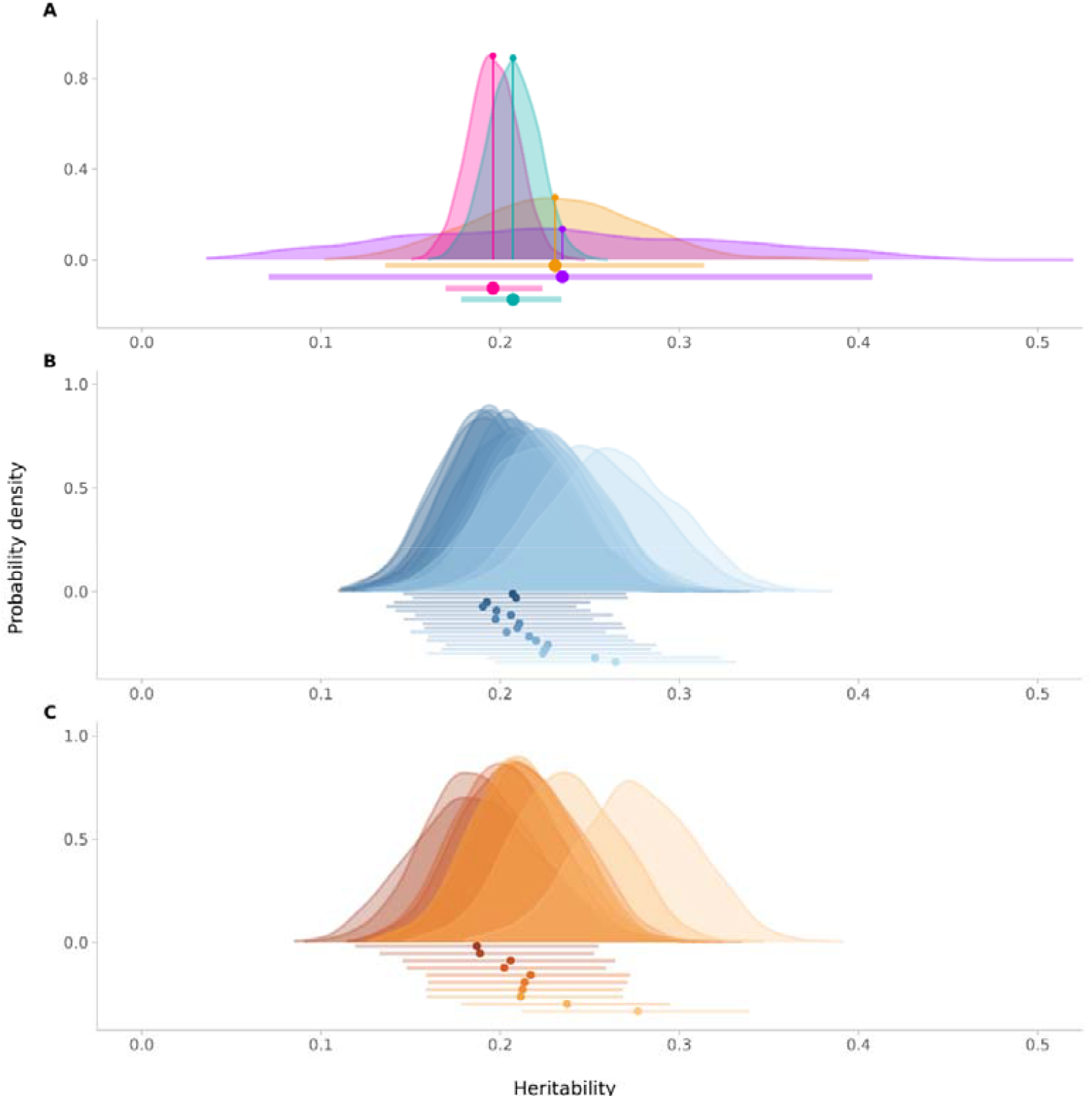
Posterior distributions of heritability estimates for birth weight by the level of systematic effects in the experimental farm dataset. A: sex; turquoise - female, pink - male, purple - hermaphrodite, yellow - sex unknown. B: litter size; darker colours indicate larger litter sizes. C: parturition number; darker colours indicate higher parturition numbers.

**Figure 4.**
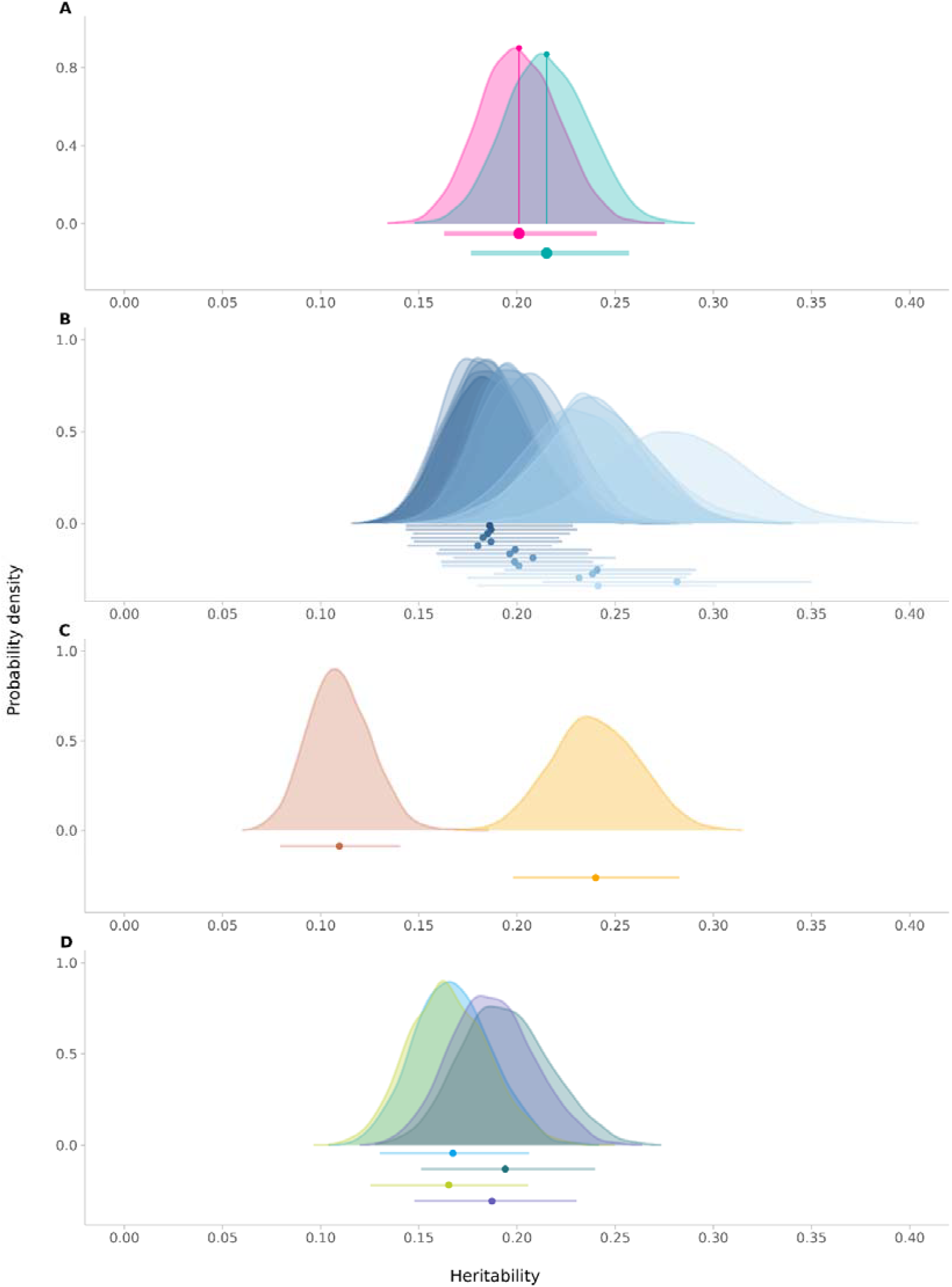
Posterior distributions of heritability estimates of birth weight by the level of systematic effects in the commercial farm dataset. A: sex; turquoise - female, pink - male. B: litter size; darker colours indicate larger litter sizes. C: sow’s age; darker colours indicate greater age. D: sires’ breed.

### Genetic correlations between means and variability

Including correlations between both litter effects, i.e., for the mean BW and its variability, improved the model’s fit to the data, but it also increased the genetic correlations between the mean BW and its variability (Table 4 and Table 5). The genetic correlation between the mean BW and its variability was always positive and ranged between 0.149 and 0.307 for the experimental farm and between 0.220 and 0.589 for the commercial farms (Figure 5). This indicates that, with selection pressure for increased BW, the variability of this trait will also increase.

**Figure 5.**
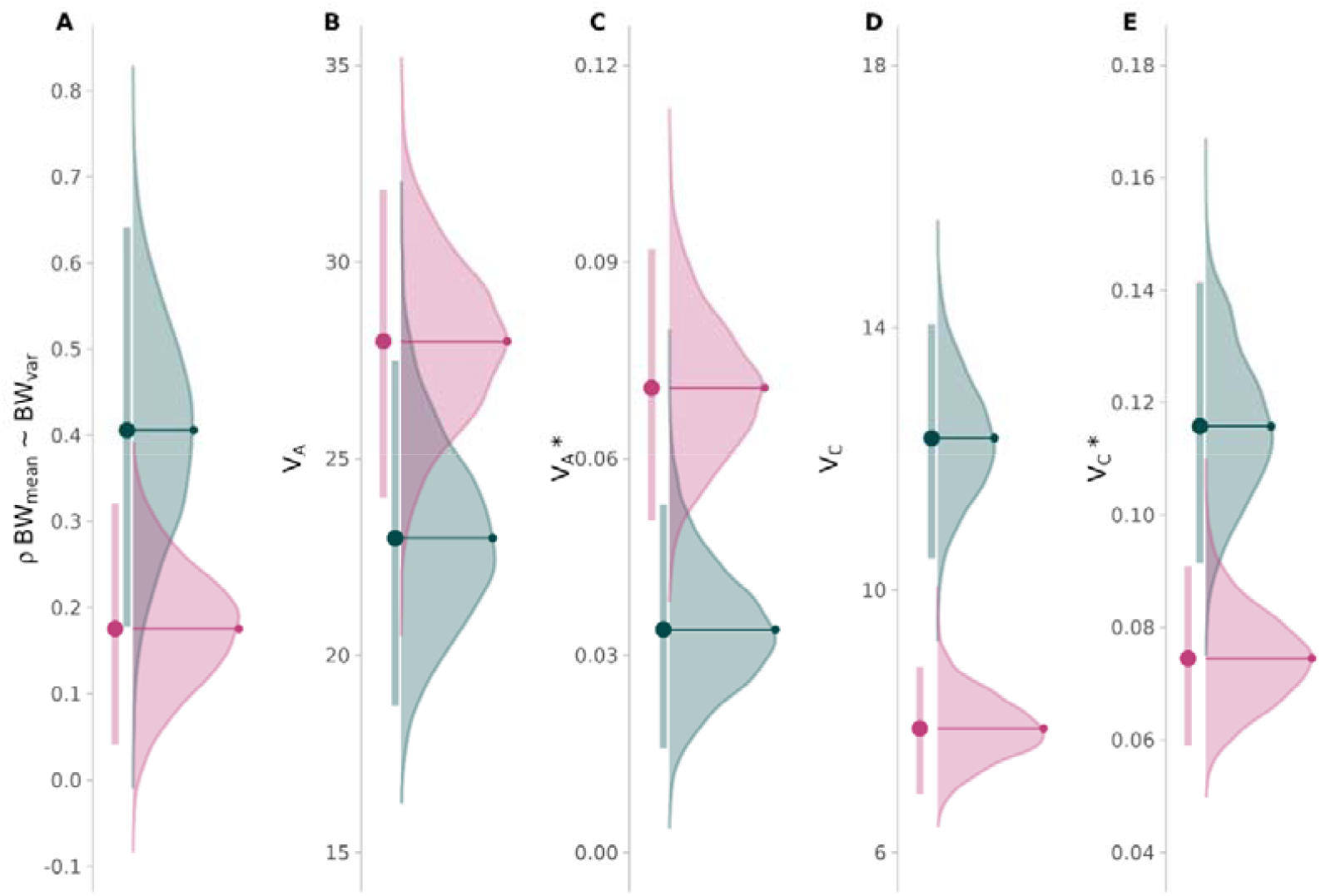
Genetic parameters of birth weight and birth weight variability on the experimental (pink) and commercial (green) farms. A: genetic correlation between the mean and the variability of birth weight, B: direct genetic effect, C: direct genetic effect of variability, D: litter effect, E: litter effect of variability. The axes of B and D are given in kg.

Negative genetic correlations were observed between the direct genetic effect on BW and the maternal effect on BW variance in both datasets and across models, except for HE_6. This finding indicates that, genetically, the increase in BW is linked to a decrease in within-mother variation in her offspring’s BW. The genetic correlations between the two genetic effects (direct and maternal) for BW were negative, except in HE_5 and HE_7 for the experimental dataset and in HE_4, HE_5, and HE_7 for the commercial dataset. However, the correlations involving the direct genetic effect on the mean and the variance level had high standard errors.

In both datasets, the best-fitting model was HE_5, which included both genetic effects for BW and the same random effects for BW and its variability and correlations between litter random effects in both datasets.

## Discussion

This study aimed to describe the genetic variation in within-litter birth weight uniformity in two populations of Swiss Large White pigs. To our knowledge, this is the first time that a heteroscedastic model including two correlated genetic effects affecting the mean of a trait has been used together with the genetic component on the residual variance.

### Importance of birth weight uniformity

Birth weight in pigs affects not only piglet survival at birth and until weaning (Moreira et al., 2020; Rutherford et al., 2013; Kapell et al., 2011; Knol et al., 2010) but also the entire production life of the pig (Beaulieu et al., 2010; Quiniou et al., 2002; Lopez-Bote, 1998). Additionally, high within-litter variation can also cause issues during birth (e.g., obstruction of the birth canal by the largest piglets), during lactation (e.g., smaller piglets have lower chances of reaching functional teats), and increase overall competition between piglets until weaning (Andersen et al., 2011; Canario et al., 2010). Uniform within-litter BW has also been linked to a reduced occurrence of intrauterine growth retardation, thereby improving piglet prenatal survival (Riddersholm et al., 2021; Matheson et al., 2018). Long-term experiments on two divergent lines of mice have shown that selection for high uniformity of within-litter BW is possible and can lead to increased survival of individuals at birth and until weaning. Additionally, uniformity was linked to greater embryonic and foetal survival, resulting in animals with greater overall robustness (Formoso-Rafferty et al., 2016, 2022). In the mouse selection experiments, the litter size at birth and weaning was greater in the line selected for increased within-litter BW uniformity than in the line selected for BW heterogeneity and compared with an unselected control line. This finding shows that by selecting for uniformity, it is possible to increase litter size without detrimental effects on BW uniformity (Formoso-Rafferty et al., 2020).

### Influence of systematic effects

The variability in birth weight increased with litter size in both datasets (lower heritabilities showing greater residual variance). Litter size has greatly increased in pig populations in recent decades through selection (Knap et al., 2023), but this has led to high variability in piglet BW within litters (Moreira et al., 2020), thus creating a competitive environment where the smallest individuals loose and die (Riddersholm et al., 2021; Damgaard et al., 2013; Quiniou et al., 2002) and leading to a reduction in the mean BW (Quesnel et al., 2008). The heterogeneity of BW increased with the number of parturitions (experimental farm) and the age of the sow (commercial farm). This has also been observed by Riddersholm et al. (Riddersholm et al., 2021), who reported that BW variability increased after the 5th parturition compared with the 2nd to 4th parturition. For commercial farms, an increase in residual variance with age was observed. With respect to sex, females are born with lower weights than males, which contributes to the heterogeneity (Wittenburg et al., 2011), but females are more homogeneous than males are, as shown by the higher heritability estimates. In the case of paternal crossbreeds, we found differences in BW means and variability; the Swiss Large White cross with service sires of other breeds had piglets with higher BW than purebred pigs did. In the literature, the crossbreed effect has been shown to be important regarding BW and its variability (Boonkum et al., 2025; Miller et al., 1979). Additionally, hyperprolific lines are more prone to have lower uniformity within-litter BW (Knap et al., 2023; Riddersholm et al., 2021); thus, increasing the litter size alone would not be advisable (Sell-Kubiak, 2021).

### Genetics of birth weight

In this study, we jointly estimated the direct and maternal genetic effects on piglet BW, which are correlated and contribute to the trait mean. In previous studies, direct heritability varied from 0.01 to 0.13, whereas maternal heritability ranged from 0.16-0.28 (Kapell et al., 2011; Knol et al., 2010; Su et al., 2008; Roehe, 1999). In our study, the inclusion of direct additive genetic variance improved the model fit, but its contribution to the total variance in BW was negligible. Similarly, paternal effects were ignored in our analysis, as previous studies indicated very low variance due to the sire effect on birth weight (Cieleń and Sell-Kubiak, 2024; Formoso-Rafferty et al., 2023). In both datasets, the additive genetic variance became insignificant when the complete model (HE_5) was fitted, even though the AIC and BIC improved. Thus, we can conclude that the additive genetic effect on birth weight is negligible compared to the maternal genetic effect on birth weight. Additionally, the correlations involving the direct effect are not different from 0. These findings allow the use of a simpler model in the future evaluation of BW.

### Genetics of birth weight uniformity

The results of our study show that the within-litter variation in BW in Swiss pigs is very similar to that reported in previous studies on other breeds of pigs (Sell-Kubiak et al., 2015; Sell-Kubiak et al., 2015; Kapell et al., 2011; Knol et al., 2010; Su et al., 2008; Roehe, 1999). Most importantly, this trait contains a genetic component that is attributed mainly to maternal additive variance, as has been shown previously in pigs (Sell-Kubiak et al., 2015) and in mice (Formoso-Rafferty et al., 2023; Pun et al., 2013).

The direct genetic effect on the variance in BW was not included for several reasons: 1) repeated records per piglet would be needed to obtain accurate estimates of variance components for variability, and 2) the literature indicates issues with including this effect (and corresponding correlations) in the heteroscedastic model (Pun et al., 2013).

As reported by us and others (Sell-Kubiak et al., 2015; Sell-Kubiak et al., 2015; Kapell et al., 2011; Knol et al., 2010; Su et al., 2008; Roehe, 1999), there is a moderate genetic correlation between the maternal additive effect on the mean and the variance in BW. Our study also revealed a difference in this correlation between commercial farms and experimental farms, which might be due to the timing of piglet weighing; if BW data are not collected 24 h after birth, BW levels are affected by postnatal conditions. However, from a genetic standpoint, the difference in the correlations between the farms might be caused by the selection for increased BW on commercial farms, affecting the magnitude of correlations between the mean and the variance.

The positive genetic correlation between BW and its variability is undesirable, as it will lead to smaller piglets if selection for decreased BW variability is implemented. However, the estimated level of correlation allows the possibility of selecting for increased mean birth weight while simultaneously selecting for greater uniformity with the appropriate selection index. This requires the simultaneous selection of both traits. On the basis of the aforementioned study in mice, selection for BW uniformity alone appears to be both practical and potentially beneficial to animal welfare. While such selection may lead to a smaller average BW, the resulting homogeneity within litters could increase survival rates until weaning. Thus, combining selection for greater within-litter uniformity with selection for greater survival at birth—a practice already implemented by breeding companies—could improve piglet survival and robustness, in line with societal demands (EFSA Panel on Animal Health and Welfare (AHAW) et al., 2022).

## Conclusions

Our results suggest that modelling BW and its variability as genetic components of environmental variance, while including only the maternal genetic effect for both traits, can yield accurate estimates of BW variability for breeding programs. This simplification is supported by the fact that the additive genetic effect plays a limited role in explaining BW variation. Additionally, despite moderate genetic correlations between the mean and the variance of BW, focusing only on the selection for uniformity in within-litter BW would be recommended rather than trying to include both traits simultaneously in the selection index.

## Declarations

### Ethical approval

The experimental procedure was approved by the Swiss Office for Food Safety and Veterinary Affairs (national licence number 36304), and all procedures were conducted in accordance with the Swiss Ordinance on Animal Protection and the Ordinance on Animal Experimentation.

### Availability of data and material

The data used in this study can be accessed upon reasonable request sent to the corresponding author.

### Author contributions

ESK – conceptualization, validation, writing – original draft, writing – review and editing

CK – conceptualization, investigation, resources, data curation, writing – review & editing, visualization

AP - investigation, resources, data curation, writing – review & editing.

JPG – conceptualization, methodology, software, validation, formal analysis, investigation, data curation, writing - original draft, writing - review & editing, visualization, supervision

NFR – conceptualization, methodology, software, validation, formal analysis, investigation, data curation, writing - original draft, writing - review & editing, visualization

NK – resources, data curation, writing - review & editing

IC – conceptualization, methodology, software, validation, formal analysis, investigation, data curation, writing - original draft, writing - review & editing, visualization, supervision

## Acknowledgements

The authors would like to thank Guy Maïkoff and his team at the experimental farm for their data collection.

